# Isolation and characterization of a novel phage SaGU1 that infects *Staphylococcus aureus* clinical isolates from patients with atopic dermatitis

**DOI:** 10.1101/2020.10.24.353185

**Authors:** Yuzuki Shimamori, Ajeng K. Pramono, Tomoe Kitao, Tohru Suzuki, Shin-ichi Aizawa, Tomoko Kubori, Hiroki Nagai, Shigeki Takeda, Hiroki Ando

## Abstract

The bacterium *Staphylococcus aureus*, which grows on healthy human skin, may cause diseases such as atopic dermatitis (AD). Treatment for such AD cases involves antibiotic use; however, alternate treatments are preferred owing to the development of antimicrobial resistance. This study aimed to characterize the novel bacteriophage SaGU1 as a potential agent for phage therapy to treat *S. aureus* infections. SaGU1 that infects *S. aureus* strains previously isolated from the skin of patients with AD was screened from sewage samples in Gifu, Japan. Its genome was sequenced and analyzed using bioinformatics tools, and the morphology, lytic activity, stability, and host range of the phage were determined. The SaGU1 genome consisted of 140,909 bp with an average GC content of 30.2%. The viral chromosome contained putative 225 protein-coding genes and four tRNA genes, carrying neither toxic nor antibiotic resistance genes. Electron microscopy analysis revealed that SaGU1 belongs to the *Myoviridae* family. Stability tests showed that SaGU1 was heat-stable under physiological and acidic conditions. Host-range testing revealed that SaGU1 could infect a broad range of *S. aureus* clinical isolates present on the skin of patients with AD, whereas it did not kill strains of *Staphylococcus epidermidis*, which are symbiotic bacteria in the human skin microbiota. Our data suggest that SaGU1 is a potential candidate for developing a phage therapy to treat AD caused by pathogenic *S. aureus*.

## Introduction

*Staphylococcus aureus* is a Gram-positive commensal bacterium present in human microbiota, but it can act as an opportunistic pathogen causing several infectious diseases, such as pneumonia, endocarditis, and bacteremia [1–4]. Atopic dermatitis (AD) is a common inflammatory skin disease that can be caused by an abnormal colonization of *S. aureus* [5, 6].

The current treatment for such AD cases includes the use of topical antibiotics; however, improvement of symptoms is temporary, and this results in the development of antibiotic resistance [7]. The growth of drug-resistant *S. aureus*, such as methicillin-resistant *S. aureus* (MRSA), may also be seen on the skin of patients with AD [8]. In a previous study in the USA, 80% of patients with AD showed colonization of *S. aureus* on the skin, 16% of which were identified as MRSA [9].

Another issue observed with using antibiotics in AD treatment involves their impact on the commensal bacterial community [10, 11]. *Staphylococcus epidermidis* is the major symbiotic bacterium present in the human skin microbiota, and providing this bacterium helps recover the skin barrier function by improving the skin microbiota in patients with AD. The lipopeptide produced by *S. epidermidis* enhances the production of antimicrobial peptides on the skin surface of humans and mice, thereby preventing infection from pathogenic bacteria including *S. aureus* [12–14]. Therefore, an efficient strategy to treat AD that does not affect the human skin microbiota is strongly desired as an alternative to antibiotic treatment.

Recently, the application of bacteriophages has gained attention as a therapeutic tool for altering the human microbiota [15, 16]. In this study, we isolated a phage SaGU1 that specifically kills *S. aureus* isolated from patients with AD. Here, we describe the genomic information, biophysical stability, and ability to infect previously identified clinical isolates of *S. aureus* and *S. epidermidis* from the skin of patients with AD.

## Materials and Methods

### Bacterial strains

All bacterial strains used in this study are listed in Table 1. *Staphylococcus* clinical isolates were obtained from Prof. Suzuki’s laboratory, Gifu University, Gifu, Japan. The bacterial species were confirmed based on the analysis of the V3-V4 region in the 16S rRNA gene [17]. All bacterial strains were grown in lysogeny broth (LB) (Formedium, UK) medium at 37 °C unless stated otherwise.

**Table 1.** Genotyping, drug resistance, and phage susceptibility of bacterial strains used in this study.

| Species | Strain | Source | MLST |  | Drug resistance | Plaque formation | EOP |
| --- | --- | --- | --- | --- | --- | --- | --- |
|  |  |  | ST | CC |  |  |  |
| *Staphylococcus aureus* | 1056-1 | Clinical isolate from an AD patient | 8 | CC8 | ABPC | + | 1.0 |
|  | 158-F1 | Clinical isolate from an AD patient | 4 | - | - | + | 0.8 |
|  | 107-1 | Clinical isolate from an AD patient | 15 | CC15 | ABPC | + | 0.8 |
|  | 159-B1 | Clinical isolate from an AD patient | 6 | CC5 | - | + | 3.3 |
|  | 163-R2 | Clinical isolate from an AD patient | 15 | CC15 | - | + | 1.7 |
|  | SA-1 | Clinical isolate from an AD patient | 8 | CC8 | ABPC, LVFX | + | 0.8 |
|  | SA-2 | Clinical isolate from an AD patient | 6 | CC5 | ABPC | + | 0.3 |
|  | SA-3 | Clinical isolate from an AD patient | 15 | CC15 | - | + | 0.8 |
|  | SA-4 | Clinical isolate from an AD patient | 8 | CC8 | - | + | 1.7 |
|  | SA-5 | Clinical isolate from an AD patient | 398 | - | - | + | 1.7 |
|  | SA-6 | Clinical isolate from an AD patient | 15 | CC15 | - | + | 0.8 |
|  | SA-7 | Clinical isolate from an AD patient | 8 | CC8 | ABPC, LVFX | + | 0.3 |
|  | SA-8 | Clinical isolate from an AD patient | 25 | - | - | - | < 10<sup>-8</sup> |
|  | GTC01187 | Clinical isolate, MRSA | 5 | CC5 | MPIPC, ABPC, CEZ, CMZ, CFX | + | 0.3 |
|  | GTC01196 | Clinical isolate, MRSA | 239 | CC8 | MPIPC, ABPC, CMZ, CFX | - | < 10<sup>-8</sup> |
|  | RN4220 | Laboratory strain | 8 | CC8 | - | + | 0.4 |
| *Staphylococcus epidermidis* | SE-1 | Clinical isolate from an AD patient | 922 | - | ABPC | - | < 10<sup>-8</sup> |
|  | SE-2 | Clinical isolate from an AD patient | 5 | - | - | - | < 10<sup>-8</sup> |
|  | SE-3 | Clinical isolate from an AD patient | 297 | - | ABPC | - | < 10<sup>-8</sup> |
|  | SE-4 | Clinical isolate from a healthy person | 152 | - | - | - | < 10<sup>-8</sup> |
|  | SE-5 | Clinical isolate from an AD patient | 2 | - | MPIPC, ABPC, CEZ, CMZ, IPM, CFX | - | < 10<sup>-8</sup> |
|  | SE-6 | Clinical isolate from a healthy person | 992 | - | - | - | < 10<sup>-8</sup> |
|  | SE-7 | Clinical isolate from an AD patient | 53 | - | - | - | < 10<sup>-8</sup> |
| *Listeria innocua* | KF2492 | Isolate from chicken | ND | ND | ND | - | < 10<sup>-8</sup> |
| *Salmonella enterica* serovar Typhimurium | LT2 | NBRC13245 | ND | ND | ND | - | < 10<sup>-8</sup> |
| *Bacillus subtilis* | NBRC3936 | NBRC3936 | ND | ND | ND | - | < 10<sup>-8</sup> |
| *Escherichia coli* | DH5α | Laboratory strain | ND | ND | ND | - | < 10<sup>-8</sup> |
| *Pseudomonas aeruginosa* | PAO1 | Laboratory strain | ND | ND | ND | - | < 10<sup>-8</sup> |
|  | PA14 | Laboratory strain | ND | ND | ND | - | < 10<sup>-8</sup> |

MLST, multilocus sequence typing; ST, sequence type; CC, clonal complex; ABPC, amoxicillin; CEZ, cefazolin sodium salt; CFX, cefoxitin; CMZ, cefmetazole sodium salt; IPM, imipenem, LVFX, levofloxacin; MPIPC, Oxacillin sodium; EOP, efficiency of plating; ND: Not determined.

### Multilocus sequence typing of *Staphylococcus* clinical isolates

Genotyping of *S. aureus* and *S. epidermidis* was performed by multilocus sequence typing (MLST) [18]. Seven housekeeping genes (*arcC*, *aroE*, *glpF*, *gmk*, *pta*, *tpi*, and *yqi* for *S. aureus*, and *arcC*, *aroE*, *gtr*, *mutS*, *pyrR*, *tpiA*, and *yqiL* for *S. epidermidis*) amplified using PCR were sequenced (Research Equipment Sharing Promotion Center NGS facility, Gifu University) and compared to the allele profiles from the database of *S. aureus* and *S. epidermidis* (https://pubmlst.org/saureus/).

### Drug susceptibility testing

Drug susceptibility was determined according to the Clinical and Laboratory Standards Institute (CLSI) guidelines [19] using a DP32 drug plate (Eiken Chemical, Japan). Briefly, the overnight cultures of the selected *Staphylococcus* isolates were diluted to an OD_600_ of approximately 0.25. Twenty-five microliters of the diluted bacterial culture were added to 12 mL of LB medium, of which 100 μL was added to each well of the DP32 drug plate. The plate was incubated at 37 °C for 16–20 h. The minimal inhibitory concentrations (MICs) of the strains were determined according to the manufacturer’s instructions and CLSI guidelines. MRSA was defined as *S. aureus* showing an MIC as follows: oxacillin (MPIPC), ≥ 4 μg/mL; cefoxitin (CFX), ≥ 8 μg/mL according to CLSI guidelines.

### Isolation and propagation of bacteriophages

Phage screening was performed using sewage samples obtained from the northern plant of the Water and Sewage Division of Gifu City, Gifu, Japan, according to the protocol published previously [20]. Briefly, 1.6 L of sewage samples were centrifuged at 8,000 × *g* for 20 min at 4 °C, and the resulting supernatant was added to polyethylene glycol 6,000 (final concentration, 10% w/v) and NaCl (final concentration, 4% w/v), and stored overnight at 4 °C. The sample was then centrifuged at 10,000 × *g* at 4 °C for 90 min, and the resulting precipitate was re-suspended in 2 mL of phage buffer (10 mM Tris-HCl [pH 7.5], 10 mM CaCl_2_, 10 mM MgSO_4_, 70 mM NaCl). After adding a few drops of chloroform, the sample was kept on ice for 6 h. The sample was then centrifuged at 8,000 × *g* for 10 min, filtrated through 0.22 μm filters (Merck), and stored at 4 °C. One hundred microliters of the solution and 200 μL of an overnight culture of each *S. aureus* strain were mixed with soft agar, then overlaid on the LB plate (double-layer plate method). After incubation at 37 °C, the obtained plaques were individually picked up and re-suspended in 100 μL of phage buffer. The titer of the phage lysate was measured by spotting 10-fold serially-diluted phages on the bacterial lawn, and calculated as plaque-forming units (PFU)/mL.

### Transmission electron microscopy (TEM)

The phage lysate was prepared for TEM according to a previously published method [21]. Samples were negatively stained with 2% phosphotungstic acid (w/v, pH 7.0) and observed using a JEM-1200EXII electron microscope (JEOL, Japan). Micrographs were taken at an accelerating voltage of 80 kV.

### One-step growth curve

A one-step growth assay was performed according to a previously published method [22]. Bacterial culture (10^6^ CFU/mL) of *S. aureus* 1056-1 was mixed with phage lysate that showed a multiplicity of infection (MOI) of 0.001, and then incubated for 10 min at 37 °C. The mixture was centrifuged at 7,000 × *g* for 10 min at 4 °C. The supernatant was discarded, and the pellet was washed twice with LB and then re-suspended in an equal volume of LB. Next, the resuspension was incubated at 37 °C at 250 rpm. Samples were collected every 10 min to measure the phage titers. The burst size was calculated using the following equation:

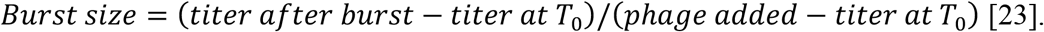

### Thermal and pH stability analysis

The thermal stability of SaGU1 was tested by incubating the phage solution in phosphate buffered saline (PBS) without calcium and magnesium [PBS (-)] at 4, 20, 30, 40, 50, 60, 70, and 80 °C for 2 h. Similarly, to assess the stability of phages under acidic and basic conditions, the phages were incubated in PBS (-) at pH 1 to 13 at 37 °C for 2 h. The rate of surviving phages was calculated using the double-layer plate method.

### Phage host range

The host range of SaGU1 was determined using the bacterial strains listed in Table 1. Two hundred microliters of stationary-phase culture of the host bacteria were mixed with 0.6% soft agar and layered on the LB plate. Then, 2.5 μL of 10-fold serially-diluted SaGU1 lysate was spotted on a plate in which the host bacteria were overlaid and incubated overnight at 37 °C. The efficiency of plating (EOP) was determined using the following equation:

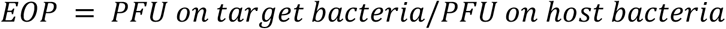

### Genome sequencing and bioinformatics analysis

The phage genome was extracted as described previously [24]. Genome sequencing was performed using the Illumina MiSeq platform (Illumina, San Diego, California, USA) and the MiSeq Reagent Kit v3 (Illumina). DNA libraries were prepared using the Nextera XT DNA Preparation Kit (Illumina) for paired-end analyses. Obtained reads were quality-filtered and assembled into contigs and scaffolds using SPAdes 3.9.0 (St. Petersburg State University, Russia) [25]. Prediction of the genes present in phage genomes was carried out using Glimmer [26] in the RAST annotation pipeline [27]. Automatic annotations were manually curated using BLASTp searches against the NCBI non-redundant protein database and NCBI Refseq viral database, with the cut-off level set to an e-value < 10^−4^. Prediction of transmembrane helices was conducted using the TMHMM Server ver. 2.0 [28, 29].

The phylogenetic trees were generated based on the JTT matrix-based model [30] of maximum likelihood method [31] using the MEGA ver. 6.0 software (Pennsylvania State University, Pennsylvania, USA) [32]. Bootstrap confidence values (500 resamplings) are as indicated on the internal branches.

### Nucleotide sequence accession number

The complete genome data of SaGU1 has been deposited in the NCBI database under accession number LC574321.

### Statistical analysis

One-step growth assay and the stability analysis data were presented as the mean ± standard deviation (SD) and analyzed using GraphPad Prism version 8.4.3 (471) (GraphPad Software, San Diego, CA, USA).

## Results and Discussion

### Isolation of phage SaGU1 infecting *S. aureus* clinical isolates

We obtained sewage samples from the northern plant of the Water and Sewage Division of Gifu City, Gifu, Japan, to screen for phages. Two clinical strains of *S. aureus* (1056-1 and 158F1) previously isolated from patients with AD were used as host bacteria to screen for the phages. In the screening process, we isolated five and three independent phages from lawns of the strains 1056-1 and 158F1, respectively. We extracted the genomic DNA from these eight phages, and determined their whole genome sequences. Unexpectedly, all of these isolates showed 100% identical genome sequences; therefore we selected a single phage infecting 1056-1 as the representative and named it SaGU1.

### Characterization of the SaGU1 genome

Sequencing of the SaGU1 genome revealed that it contained a double-stranded DNA genome of 140,909 bp in size, and an overall GC content of 30.2% (Fig. 1a). Based on its genome size and sequence comparison, SaGU1 was found to belong to the Class III *Staphylococcus* phages [33]. The SaGU1 genome encodes a total of 225 putative open reading frames (ORFs) and four transfer RNAs (tRNAs). The coding density of the genome was found to be 91%, leaving a very small intergenic region, with a mean ORF length of 567 nucleotides. All of the discovered ORFs were used against the Refseq protein database, however, only 70 ORFs had been assigned to the functions (Supplementary Table S1).

**Fig. 1.**
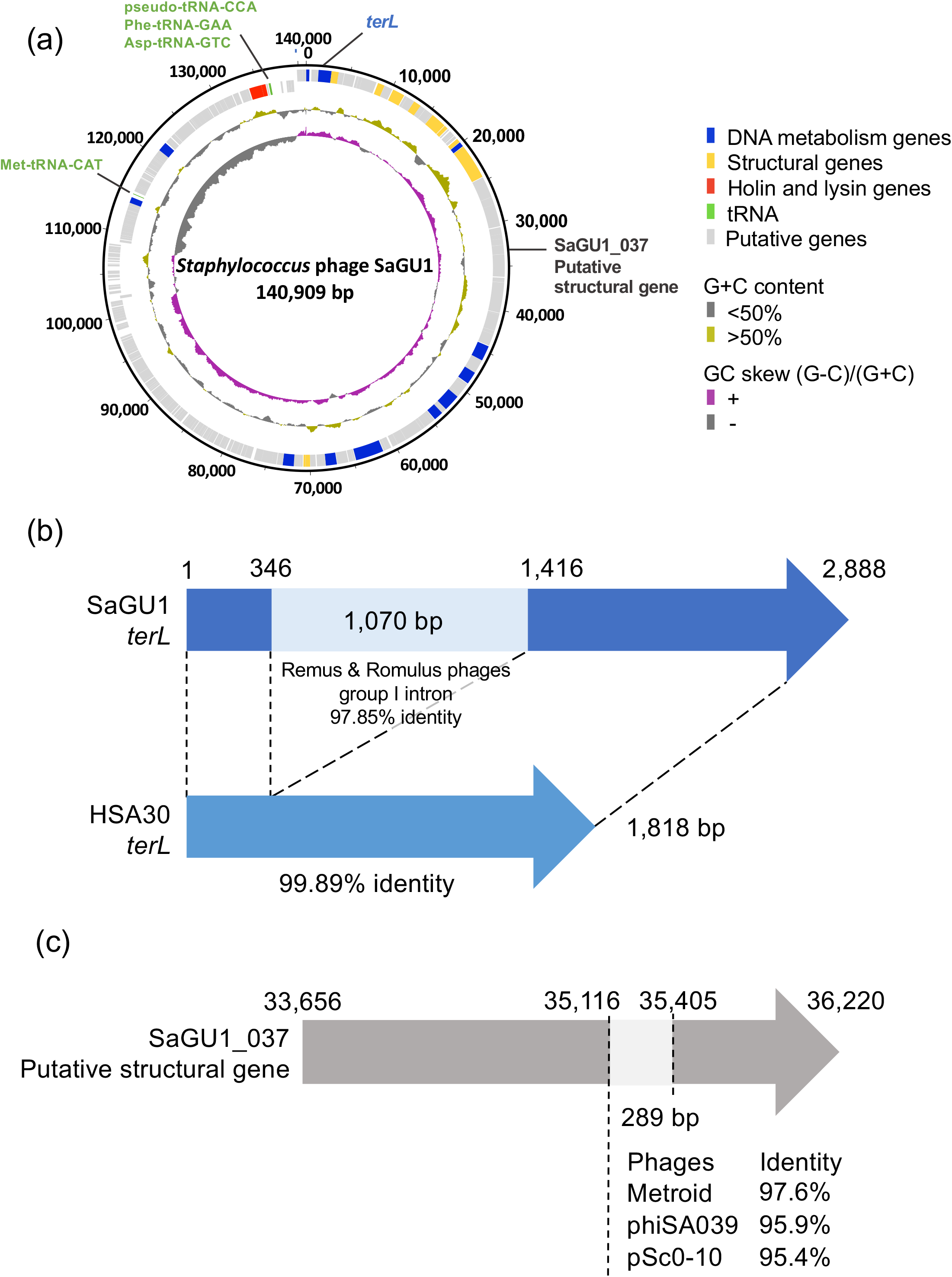
Genomic structure of the phage SaGU1. (a) Circular representation of the SaGU1 genome. Concentric rings denote the following features (from the outer to inner rings): nucleotide positions are forward strand (outer) and reverse strand (inner); the predicted genes are DNA metabolism genes (blue), structural genes (yellow), holin and lysin genes (red), tRNA (green), and putative genes (light gray); G+C content is < 50% (gray), > 50% (gold); GC skew is (G-C)/(G+C)(gray, −; purple, +). (b) Structure of SaGU1 *terL*. Pale blue line represents the region with high sequence similarity to the group I intron of *Staphylococcus* phages Remus and Romulus. (c) Structure of the SaGU1_037 putative structural gene. Pale gray line represents the region with high sequence similarity to sequences of *Staphylococcus* phages Metroid, phiSA039, and pSc0-10.

Topologically, the genome of SaGU1 can be divided into two unequal regions; the majority of the ORFs were located on the forward strand, whereas all of the tRNAs were located on the reverse strand. Met-tRNA-CAT between SaGU1_178 and SaGU1_179, and pseudo-tRNA-CCA, Phe-tRNA-GAA, and Asp-tRNA-GTC between SaGU1_218 and SaGU1_219 were also separated into two regions. More interestingly, the tRNA gene sequences were identical; this arrangement was conserved in other similar *Staphylococcus* phages. The presence of tRNA in phage genomes is not uncommon, since it has been hypothesized that viral tRNA compensates for the difference in codon usage bias between a phage and its bacterial host, and that the tRNAs correspond to codons that may be inefficiently translated by the host translational machinery [34].

The SaGU1 genome was almost identical to that of *S. aureus* phage HSA30 (MG557618); however, it contained two unique regions that the genome of HSA30 does not contain. The first region was a locus from 1 to 2,888 (Fig. 1b, Table S1). It contained an inserted sequence that spliced the *terL* gene into two parts (SaGU1_1 and SaGU1_3) and was 99.89% identical to the group I intron of Class III *Staphylococcus* phages, Remus (NC_022090) and Romulus (NC_020877) [35]. Introns are sometimes discovered in phage genomes [36, 37]. The intron of some Twort-like phages has the ability of self-splicing from RNA transcripts [37, 38], thus SaGU1_1 and SaGU1_3 are probably self-spliced to express the functional TerL [35, 39]. However, based on the BLASTn search for the NT database, the intron found in the SaGU1 genome was only present in five other genomes of Class III *Staphylococcus* phages, namely Remus, Romulus, MCE-2014 (NC_025416), StAP1 (KX532239), and phiIPLA-RODI (NC_028765) [35, 40, 41]. The other unique region compared with the genome of HSA30 was a locus from 35,116 to 35,405 in SaGU1_037 (Fig. 1c). It showed 98%, 96%, and 95% nucleotide similarity with a region on the genomes of *Staphylococcus* phage Metroid (MT411892), phiSA039 (AP018375), and pSc0-10 (KX011028), respectively. These mosaicisms indicate that horizontal gene transfer is common within this phage group, as mentioned in earlier studies [33, 42–44].

Mosaicism of phage genomes is common, but the head, tail, DNA replication, and nucleotide metabolism genes “travel together through evolution” [45], with little or no evidence of horizontal swapping. Phylogenetic analysis was also performed based on the terminase large subunit TerL, except for the intron and baseplate amino acid sequences. The TerL phylogenetic tree showed that the phage SaGU1 was clustered with *Staphylococcus* phages HSA30 and GH15 (NC_019448), showing a high bootstrap value (90%) (Fig. 2a). The baseplate tree showed that SaGU1 was clustered with the same group of *Staphylococcus* phages, GH15, JD007 (NC_019726), and G1 (NC_007066) (Fig. 2b).The phage Remus, which shared a similar intron sequence to that of SaGU1, was placed in a separate clade on the tree. This suggests that the intron was most likely obtained by a horizontal transfer.

**Fig. 2.**
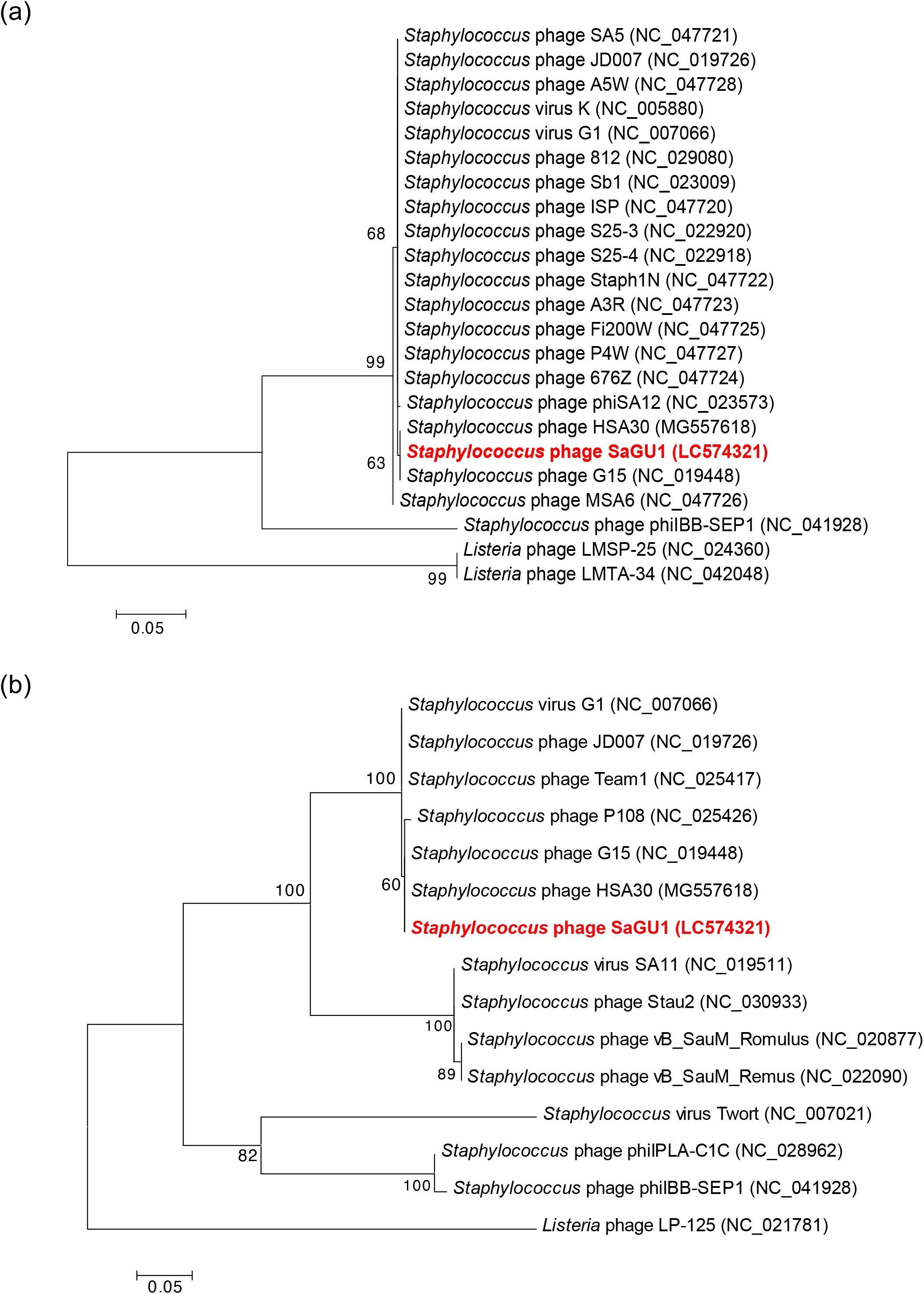
Phylogenic analysis of *Staphylococcus* phage SaGU1. Bootstrap confidence values (500 resamplings) are indicated on the internal branches. (a) Maximum likelihood trees of TerL subunit based on deduced 594 amino acids. The *Listeria* phages LMTA-34 and LMSP-25 were used as the outgroups. The phylogenetic trees were created with the exception of the intron. (b) Maximum likelihood trees of baseplate proteins based on deduced 348 amino acids. The putative baseplate assembly protein of the *Listeria* phage LP-125 was used as the outgroup.

The ORFs with predicted functions consisted of modules for virion structure, nucleotide replication and metabolism, and lysis. The structural modules were mainly located between bases 822 – 44,232. However, two ORFs for major tail proteins (SaGU1_71 and SaGU1_72) were located approximately 30 kb upstream from other structural genes, similar to members of Kayviruses, such as GH15, MCE-2014, and phILA-RODI [40, 41, 46]. The SaGU1 genome possessed ORFs encoding structural proteins typically present in members of *Myoviridae*, for example, tail sheath protein (SaGU1_20), tail tube protein (SaGU1_21), and baseplate protein (SaGU1_36) which controls its contractile tails [47].

The lysis module of SaGU1 consisted of two adjacent genes, lysin (SaGU1_216) and holin (SaGU1_217). Based on *in silico* prediction, the holin identified was a member of class II holins, which consist of two transmembrane helical domains, with both the N- and C-termini present in the cytoplasm [48]. This module was located downstream of the possible pseudo-tRNA, Phe-tRNA, and Asp-tRNA sequences. As there were no lysogeny-related genes found in the genome, phage SaGU1 most likely depends on the lytic cycle to replicate.

### Morphology of SaGU1

The TEM analysis confirmed that SaGU1 possessed an icosahedral head with a diameter of 86.7 ± 5.0 nm (*n* = 3), and a contractile tail with a length of 222.7 ± 1.9 nm (*n* = 3) and a width of 19.3 ± 0.7 nm (*n* = 3) (Fig. 3a, b). The myovirus tail is concentric with a tail tube inside a tail sheath. The contracted tail sheath is 95.0 ± 5.0 nm (*n* = 3) in length, about 42.7% of the length before contraction and a 72.5 ± 2.5 nm (*n* = 3) tail tube protrudes from under the base plate. A particle with a black head is an empty particle with no DNA (Fig. 3b). When a myovirus infects a host bacterium, the tail contracts and the DNA in the head comes out. SaGU1 has a double base plate (Fig. 3b) and no tail fibers but globular structures at the tail tip (Fig. 3a). The shape of whole virus resembled the morphology of other staphylococcal phages, Team1 (NC_025417) [49] and Remus [35]. These morphological characteristics show that the phage SaGU1 belongs to the genus Twort-like phages of the family *Myoviridae* [50, 51].

**Fig. 3.**
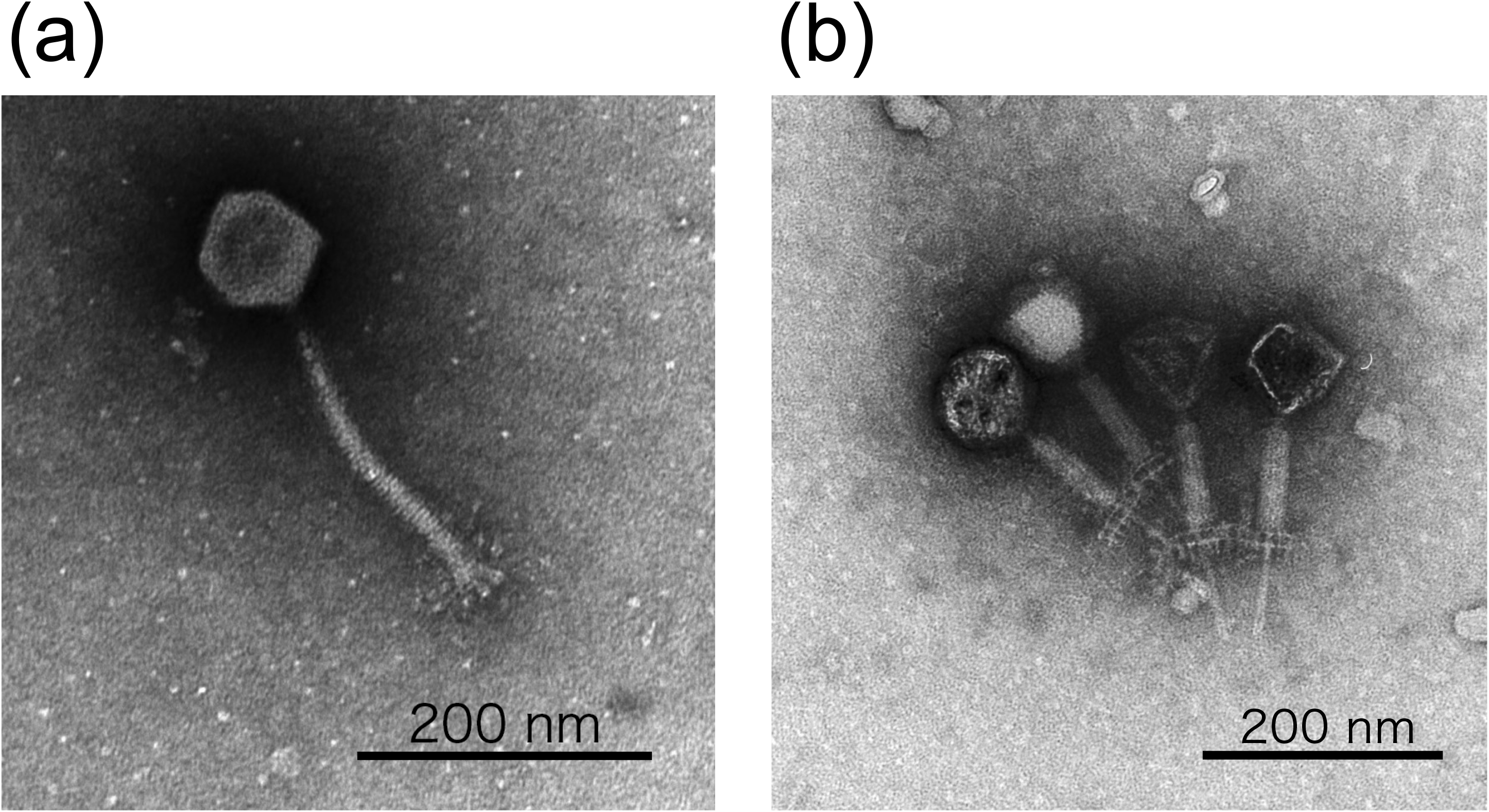
The electron micrographs of phage SaGU1. Pictures were taken by transmission electron microscopy. (a) SaGU1 normal particle. (b) SaGU1 particle with the tail contracted. A characteristic feature of the *Myoviridae* family involves the contraction of the tail sheath, and protrusion of the tail tube from the tip of the tail. A particle with a black head is an empty particle with no DNA. Scale bar, 200 nm.

### Life cycle of SaGU1

A one-step growth experiment was performed to analyze the life cycle of SaGU1 (Fig. 4a). The latent phase was found to be 40 min, followed by a growth phase lasting for 50 min. A growth plateau was reached within 90 min. The burst size of SaGU1 was calculated as approximately 26 PFU/cell. The life cycle of SaGU1 was comparable to those of other *Staphylococcus* phages in the *Myoviridae* family, phiIPLA-RODI (25 PFU/cell), phiIPLA-C1C (NC_028962) (15 PFU/cell), and Team 1 (31 PFU/cell) [41, 49].

**Fig. 4.**
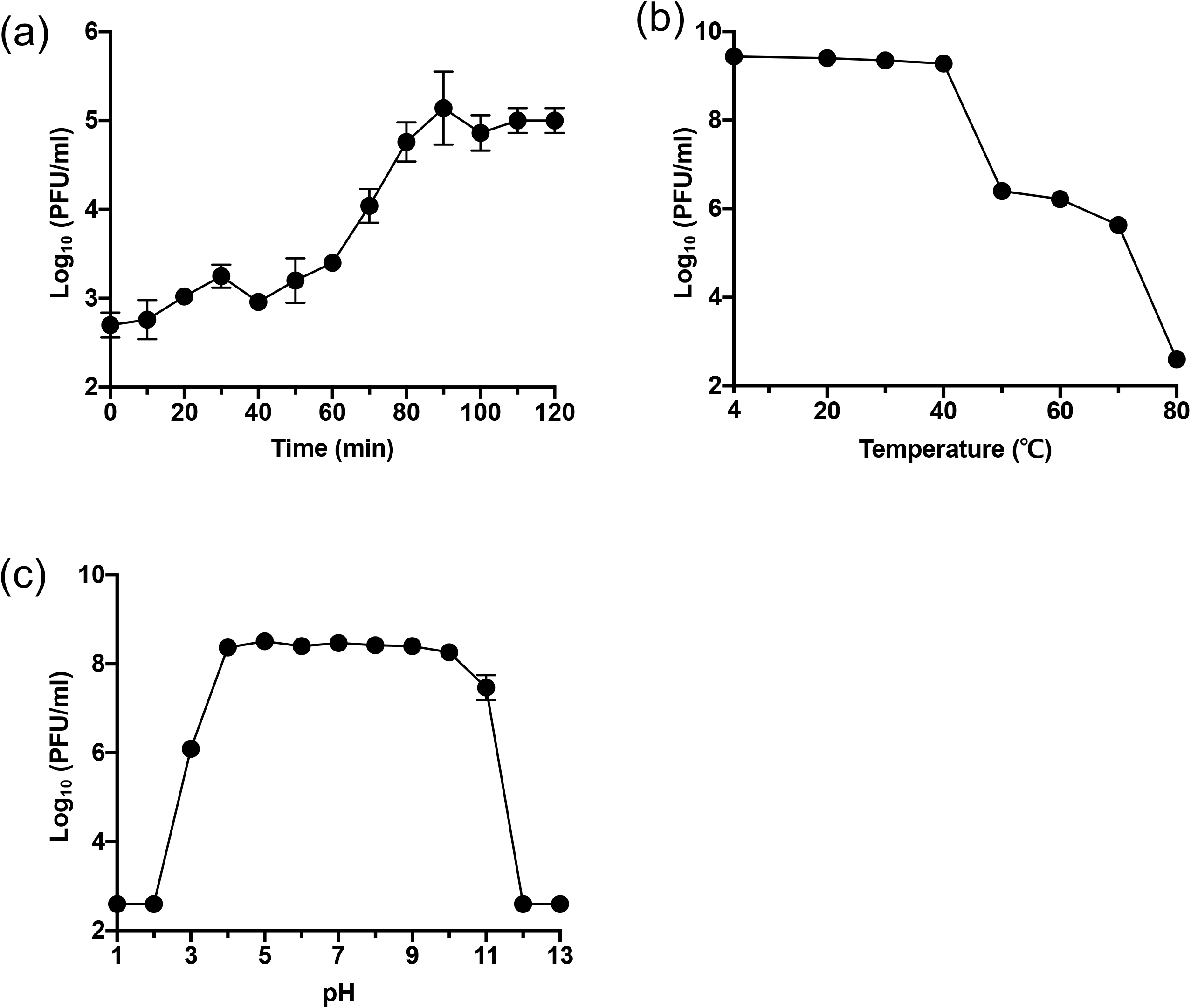
Characterization of phage SaGU1. (a) One-step growth curve of SaGU1 at 37 °C. (b) Effect of temperature on the lytic activity of SaGU1. (c) Effect of pH on the lytic activity of SaGU1. The data are presented as the mean ± standard deviation of at least three independent experiments. Small error bars are obscured by symbols. The detection limit was 4.0 × 10^2^ PFU/mL.

### Biophysical stability of SaGU1

Stable phages are required for phage therapy [52]. We examined whether SaGU1 was stable at various temperatures and pH values. Results of thermostability tests indicated that SaGU1 was stable at temperatures ranging from 4 °C to 40 °C. The stability gradually decreased between 50 and 70 °C, and the phage was inactivated at 80 °C (Fig. 4b). The effects of high and low pH on the stability of SaGU1 were also examined (Fig. 4c). SaGU1 showed stable lytic activity after incubation at conditions between pH 4 and pH 10, whereas it drastically lost lytic activity after incubation at conditions below pH 2 and above pH 12. The human skin has a pH of approximately 4.1 to 5.8, with a slightly higher pH in the skin of patients with AD (approximately pH 5.5). A pH of 7.5 is optimal for *S. aureus* growth [53, 54]. These results show that SaGU1 application will be stable on the skin of patients with AD, making it potentially useful for phage therapy.

### Host specificity of SaGU1

The host specificity of SaGU1 was examined using *Staphylococcus* strains listed in Table 1. The EOP assay results showed that SaGU1 could infect 14 out of 16 *S. aureus* strains (EOP values ≥ 0.3). The results of MLST analysis indicated that SaGU1 infected *aureus* strains that belong to diverse genomic lineages of sequence type (ST) 4, ST5, ST8, ST6, ST15, and ST398 (*S. aureus* 1056-1 and 158-F1 used for the screening of SaGU1, belonged to ST8 and ST4, respectively), suggesting that SaGU1 infects a broad host range of *S. aureus*. Notably, SaGU1 infected the strain GTC01187 (ST5, CC5) that shows resistance to multiple antibiotics, including methicillin. However, SaGU1 did not infect *S. aureus* SA-8 and GTC01196, which belonged to ST25 and ST239, respectively.

We additionally obtained a total of seven known clinical isolates of *S. epidermidis* (two strains from healthy people and five strains from patients with AD), which belong to different genomic lineages based on the MLST analysis, to assess whether SaGU1 infects *S. epidermidis*. The results of EOP assays showed that SaGU1 did not infect any of the strains of *S. epidermidis* (EOP >10^8^). Additionally, SaGU1 did not infect any other Gram-positive and Gram-negative bacteria, as described in Table 1.

Taken together, these data indicate that SaGU1 is a staphylococcal phage that specifically infected *S. aureus* strains of different STs but not *S. epidermidis*. It will be useful to establish the host determinants of SaGU1 in the future.

## Conclusions

A novel *Staphylococcus* phage SaGU1, which was stable under specific physiological and acidic conditions, was identified and its complete genomic sequence was determined in this study. Given that SaGU1 can specifically infect *S. aureus* from patients with AD, but not *S. epidermidis*, it may be a strong candidate for developing phage therapy to treat AD.

## Acknowledgements

This work was supported by the Japan Society for the Promotion of Science (JSPS), KAKENHI Grant Number JP15K21770 to H.A., and S.T. was awarded the Gunma University Medical Innovation Project grant. *Salmonella enterica* serovar Typhimurium LT2 (NBRC13245)*, and Bacillus subtilis* NBRC3926 (NBRC3936) strains were provided by the National Institute of Technology and Evaluation (NITE) Biological Resource Center (NBRC). *Pseudomonas aeruginosa* PAO1 and PA14 strains were kindly provided by Laurence G. Rahme (Harvard Medical School). *Staphylococcus aureus* RN4220 strain was kindly provided by Longzhu Cui and Kotaro Kiga (Jichi Medical University, Japan). *Listeria innocua* KF2492 strain was kindly provided by Noriko Nakanishi and Tomotada Iwamoto (Kobe City Environmental Health Research Institute).

## Declarations

### Funding

This work was supported by the Japan Society for the Promotion of Science (JSPS), KAKENHI Grant Number JP15K21770 to H.A. and Gunma University Medical Innovation Project grant to S.T.

### Conflicts of interest/Competing interests

The authors declare that they have no conflicts of interest.

### Ethics approval

Not applicable.

### Consent to participate

Not applicable.

### Consent for publication

The authors grant the publisher consent to publish the study.

### Code availability

Not applicable

## Author Contributions

S.T. and H.A. conceived the study. Y.S., A.K.P., T. Kitao, and S.A. performed experiments. T.S. contributed to obtaining clinical isolates. Y.S., A.K.P., T. Kitao, T.S., S. Kubori, H.N., S.T., and H.A. analyzed the data. Y.S., A.K.P., T. Kitao, S.T., and H.A. wrote the manuscript. All authors have read and approved the final version of the manuscript.

## Notes

### Competing Interest Statement

The authors have declared no competing interest.

