## Supplementary Table S1 for "Isolation and characterization of a novel phage SaGU1 that infects *Staphylococcus aureus* clinical isolates from patients with atopic dermatitis"

Table S1. Annotated gene list of phage SaGU1

| Locus tag | Type | Strand | Start | Stop | Predicted Function |
|---|---|---|---|---|---|
| SaGU1_001 | CDS | + | 1 | 354 | Phage terminase, large subunit |
| SaGU1_002 | CDS | + | 591 | 1361 | Hypothetical protein |
| SaGU1_003 | CDS | + | 1401 | 2888 | Phage terminase, large subunit |
| SaGU1_004 | CDS | + | 2881 | 3702 | Putative structural head protein |
| SaGU1_005 | CDS | + | 3689 | 3862 | Hypothetical protein |
| SaGU1_006 | CDS | + | 3859 | 4338 | Hypothetical protein |
| SaGU1_007 | CDS | + | 4431 | 5609 | Putative membrane protein |
| SaGU1_008 | CDS | + | 5751 | 6035 | Putative membrane protein |
| SaGU1_009 | CDS | + | 6053 | 6424 | Putative portal protein |
| SaGU1_010 | CDS | + | 6428 | 8119 | Putative portal protein |
| SaGU1_011 | CDS | + | 8312 | 9085 | Putative prohead protease |
| SaGU1_012 | CDS | + | 9104 | 10063 | Hypothetical protein |
| SaGU1_013 | CDS | + | 10179 | 11570 | Phage major capsid protein |
| SaGU1_014 | CDS | + | 11662 | 11958 | Hypothetical protein |
| SaGU1_015 | CDS | + | 11971 | 12879 | Hypothetical protein |
| SaGU1_016 | CDS | + | 12893 | 13771 | Capsid protein |
| SaGU1_017 | CDS | + | 13771 | 14391 | Hypothetical protein |
| SaGU1_018 | CDS | + | 14410 | 15246 | Hypothetical protein |
| SaGU1_019 | CDS | + | 15248 | 15463 | Hypothetical protein |
| SaGU1_020 | CDS | + | 15490 | 17253 | Tail sheath protein |
| SaGU1_021 | CDS | + | 17326 | 17754 | Tail tube protein |
| SaGU1_022 | CDS | + | 17844 | 18002 | Hypothetical protein |
| SaGU1_023 | CDS | + | 17992 | 18132 | Hypothetical protein |
| SaGU1_024 | CDS | + | 18175 | 18627 | Hypothetical protein |
| SaGU1_025 | CDS | + | 18640 | 18834 | Hypothetical protein |
| SaGU1_026 | CDS | + | 18905 | 19216 | Hypothetical protein |
| SaGU1_027 | CDS | + | 19348 | 19806 | Phage tail tape measure |
| SaGU1_028 | CDS | + | 19850 | 20386 | RNA polymerase |
| SaGU1_029 | CDS | + | 20442 | 24497 | Phage tail tape measure |
| SaGU1_030 | CDS | + | 24575 | 27001 | Secretory antigen SsaA-like protein |
| SaGU1_031 | CDS | + | 27015 | 27902 | Hypothetical protein |
| SaGU1_032 | CDS | + | 27902 | 30448 | Glycerophosphoryl diester phosphodiesterase |
| SaGU1_033 | CDS | + | 30555 | 31346 | Hypothetical protein |
| SaGU1_034 | CDS | + | 31346 | 31870 | Hypothetical protein |
| SaGU1_035 | CDS | + | 31870 | 32574 | Hypothetical protein |
| SaGU1_036 | CDS | + | 32589 | 33635 | Baseplate protein |
| SaGU1_037 | CDS | + | 33656 | 36220 | Putative structural protein |
| SaGU1_038 | CDS | + | 36331 | 36852 | Hypothetical protein |
| SaGU1_039 | CDS | + | 36873 | 40331 | Adsorption-associated tail protein |
| SaGU1_040 | CDS | + | 40380 | 40538 | Hypothetical protein |
| SaGU1_041 | CDS | + | 40539 | 42461 | Hypothetical protein |
| SaGU1_042 | CDS | + | 42475 | 42849 | Hypothetical protein |
| SaGU1_043 | CDS | + | 42856 | 44232 | Phage capsid and scaffold |
| SaGU1_044 | CDS | + | 44322 | 46070 | Phage DNA helicase |
| SaGU1_045 | CDS | + | 46082 | 47695 | Putative Rep protein |
| SaGU1_046 | CDS | + | 47688 | 49130 | Phage DNA helicase |
| SaGU1_047 | CDS | + | 49209 | 49628 | Hypothetical protein |
| SaGU1_048 | CDS | + | 49628 | 50653 | Phage recombination exonuclease |
| SaGU1_049 | CDS | + | 50653 | 51030 | Hypothetical protein |
| SaGU1_050 | CDS | + | 51030 | 52949 | Phage recombination related exonuclease |
| SaGU1_051 | CDS | + | 52949 | 53545 | Hypothetical protein |
| SaGU1_052 | CDS | + | 53560 | 54627 | Phage DNA primase/helicase |
| SaGU1_053 | CDS | + | 54692 | 55030 | Hypothetical protein |
| SaGU1_054 | CDS | + | 55030 | 55482 | Hypothetical protein |
| SaGU1_055 | CDS | + | 55469 | 56077 | Hypothetical protein |
| SaGU1_056 | CDS | + | 56095 | 56487 | Ribonucleotide reduction protein NrdI |
| SaGU1_057 | CDS | + | 56502 | 58616 | Ribonucleotide reductase of class Ib (aerobic), alpha subunit (EC 1.17.4.1) |
| SaGU1_058 | CDS | + | 58630 | 59679 | Ribonucleotide reductase of class Ib (aerobic), beta subunit (EC 1.17.4.1) |
| SaGU1_059 | CDS | + | 59697 | 60026 | Hypothetical protein |
| SaGU1_060 | CDS | + | 60010 | 60330 | Phage oxidoreductase |
| SaGU1_061 | CDS | + | 60537 | 61133 | Hypothetical protein |

| | | | | | |
|---|---|---|---|---|---|
| SaGU1_062 | CDS | + | 61143 | 61448 | Phage integration host factor |
| SaGU1_063 | CDS | + | 61524 | 64742 | DNA polymerase I |
| SaGU1_064 | CDS | + | 64812 | 65054 | Hypothetical protein |
| SaGU1_065 | CDS | + | 65071 | 65553 | Hypothetical protein |
| SaGU1_066 | CDS | + | 65640 | 66911 | Hypothetical protein |
| SaGU1_067 | CDS | + | 66971 | 68227 | Phage recombinase |
| SaGU1_068 | CDS | + | 68231 | 68584 | Hypothetical protein |
| SaGU1_069 | CDS | + | 68571 | 69233 | Putative sigma factor |
| SaGU1_070 | CDS | + | 69361 | 69993 | Putative Ig-like protein |
| SaGU1_071 | CDS | + | 70015 | 70527 | Phage major tail protein |
| SaGU1_072 | CDS | + | 70542 | 70769 | Phage major tail protein |
| SaGU1_073 | CDS | + | 70864 | 71124 | Hypothetical protein |
| SaGU1_074 | CDS | + | 71128 | 71883 | Hypothetical protein |
| SaGU1_075 | CDS | + | 71876 | 73126 | Putative DNA repair exonuclease |
| SaGU1_076 | CDS | + | 73140 | 73508 | Hypothetical protein |
| SaGU1_077 | CDS | + | 73495 | 73806 | Hypothetical protein |
| SaGU1_078 | CDS | + | 73870 | 74406 | Hypothetical protein |
| SaGU1_079 | CDS | + | 74399 | 75166 | Hypothetical protein |
| SaGU1_080 | CDS | + | 75144 | 75590 | Hypothetical protein |
| SaGU1_081 | CDS | + | 75590 | 76453 | Hypothetical protein |
| SaGU1_082 | CDS | + | 76825 | 77556 | Hypothetical protein |
| SaGU1_083 | CDS | + | 77574 | 78032 | Hypothetical protein |
| SaGU1_084 | CDS | + | 78097 | 78540 | Hypothetical protein |
| SaGU1_085 | CDS | + | 78557 | 79261 | Hypothetical protein |
| SaGU1_086 | CDS | + | 79323 | 79721 | Putative membrane protein |
| SaGU1_087 | CDS | + | 79868 | 80110 | Hypothetical protein |
| SaGU1_088 | CDS | + | 80115 | 80672 | Hypothetical protein |
| SaGU1_089 | CDS | + | 80708 | 80884 | Hypothetical protein |
| SaGU1_090 | CDS | + | 80874 | 81125 | Hypothetical protein |
| SaGU1_091 | CDS | + | 81118 | 81351 | Hypothetical protein |
| SaGU1_092 | CDS | + | 81432 | 82076 | Putative ribulose-1,5-bisphosphate carboxylase/oxygenase small subunit |
| SaGU1_093 | CDS | + | 82092 | 82340 | Hypothetical protein |
| SaGU1_094 | CDS | + | 82352 | 82528 | Hypothetical protein |
| SaGU1_095 | CDS | + | 82521 | 82817 | Hypothetical protein |
| SaGU1_096 | CDS | + | 82865 | 83047 | Putative membrane protein |
| SaGU1_097 | CDS | + | 83060 | 83428 | Hypothetical protein |
| SaGU1_098 | CDS | + | 83441 | 83788 | Hypothetical protein |
| SaGU1_099 | CDS | + | 83788 | 84066 | Putative membrane protein |
| SaGU1_100 | CDS | + | 84136 | 84441 | Hypothetical protein |
| SaGU1_101 | CDS | + | 84456 | 84806 | Hypothetical protein |
| SaGU1_102 | CDS | + | 84806 | 85408 | Hypothetical protein |
| SaGU1_103 | CDS | + | 85422 | 85601 | Hypothetical protein |
| SaGU1_104 | CDS | + | 85827 | 86237 | Putative membrane protein |
| SaGU1_105 | CDS | + | 86239 | 86532 | Hypothetical protein |
| SaGU1_106 | CDS | + | 86549 | 86836 | Putative membrane protein |
| SaGU1_107 | CDS | + | 86847 | 86960 | Hypothetical protein |
| SaGU1_108 | CDS | + | 86953 | 87222 | Hypothetical protein |
| SaGU1_109 | CDS | + | 87298 | 87603 | Hypothetical protein |
| SaGU1_110 | CDS | + | 87603 | 88010 | Hypothetical protein |
| SaGU1_111 | CDS | + | 88021 | 88257 | Hypothetical protein |
| SaGU1_112 | CDS | + | 88254 | 88781 | Phosphoesterase |
| SaGU1_113 | CDS | + | 88762 | 89073 | Hypothetical protein |
| SaGU1_114 | CDS | + | 89119 | 89298 | Hypothetical protein |
| SaGU1_115 | CDS | + | 89313 | 89576 | Hypothetical protein |
| SaGU1_116 | CDS | + | 89579 | 89896 | Hypothetical protein |
| SaGU1_117 | CDS | + | 89897 | 90550 | Hypothetical protein |
| SaGU1_118 | CDS | + | 90628 | 90831 | Hypothetical protein |
| SaGU1_119 | CDS | + | 90847 | 91005 | Putative membrane protein |
| SaGU1_120 | CDS | + | 91021 | 91245 | Hypothetical protein |
| SaGU1_121 | CDS | + | 91258 | 91458 | Hypothetical protein |
| SaGU1_122 | CDS | + | 91459 | 91749 | Putative membrane protein |
| SaGU1_123 | CDS | + | 91842 | 92135 | Hypothetical protein |
| SaGU1_124 | CDS | + | 92132 | 93040 | Phage ribose-phosphate pyrophosphokinase |

| | | | | | |
|---|---|---|---|---|---|
| SaGU1_125 | CDS | + | 93058 | 94527 | Nicotinamide phosphoribosyltransferase |
| SaGU1_126 | CDS | + | 94606 | 94851 | Hypothetical protein |
| SaGU1_127 | CDS | + | 94871 | 95263 | Hypothetical protein |
| SaGU1_128 | CDS | + | 95265 | 95462 | Hypothetical protein |
| SaGU1_129 | CDS | + | 95528 | 95839 | Hypothetical protein |
| SaGU1_130 | CDS | + | 95845 | 96144 | Hypothetical protein |
| SaGU1_131 | CDS | + | 96144 | 96383 | Hypothetical protein |
| SaGU1_132 | CDS | + | 96373 | 96534 | Hypothetical protein |
| SaGU1_133 | CDS | + | 96531 | 96866 | Hypothetical protein |
| SaGU1_134 | CDS | + | 96866 | 97084 | Hypothetical protein |
| SaGU1_135 | CDS | + | 97122 | 97397 | Hypothetical protein |
| SaGU1_136 | CDS | + | 97459 | 97626 | Hypothetical protein |
| SaGU1_137 | CDS | + | 97663 | 97764 | Hypothetical protein |
| SaGU1_138 | CDS | + | 98495 | 98791 | Hypothetical protein |
| SaGU1_139 | CDS | + | 98803 | 98973 | Hypothetical protein |
| SaGU1_140 | CDS | + | 98987 | 99172 | Hypothetical protein |
| SaGU1_141 | CDS | + | 99441 | 99755 | Hypothetical protein |
| SaGU1_142 | CDS | + | 99768 | 100058 | Terminal repeat-encoded protein |
| SaGU1_143 | CDS | + | 100058 | 100345 | Terminal repeat-encoded protein |
| SaGU1_144 | CDS | + | 100345 | 100638 | Terminal repeat-encoded protein |
| SaGU1_145 | CDS | + | 100642 | 100890 | Terminal repeat-encoded protein |
| SaGU1_146 | CDS | + | 100903 | 101142 | Terminal repeat-encoded protein |
| SaGU1_147 | CDS | + | 101254 | 101493 | Hypothetical protein |
| SaGU1_148 | CDS | + | 101505 | 101849 | Terminal repeat-encoded protein |
| SaGU1_149 | CDS | - | 102388 | 102050 | Hypothetical protein |
| SaGU1_150 | CDS | + | 102701 | 103009 | Hypothetical protein |
| SaGU1_151 | CDS | + | 103215 | 103502 | Hypothetical protein |
| SaGU1_152 | CDS | + | 103552 | 103743 | Hypothetical protein |
| SaGU1_153 | CDS | + | 104261 | 104419 | Hypothetical protein |
| SaGU1_154 | CDS | + | 104586 | 104909 | Hypothetical protein |
| SaGU1_155 | CDS | + | 105009 | 105389 | Hypothetical protein |
| SaGU1_156 | CDS | + | 105881 | 106102 | Hypothetical protein |
| SaGU1_157 | CDS | + | 106183 | 106320 | Hypothetical protein |
| SaGU1_158 | CDS | + | 106391 | 106555 | Hypothetical protein |
| SaGU1_159 | CDS | + | 106646 | 106882 | Hypothetical protein |
| SaGU1_160 | CDS | + | 106962 | 107432 | Hypothetical protein |
| SaGU1_161 | CDS | + | 107501 | 107764 | Hypothetical protein |
| SaGU1_162 | CDS | + | 107768 | 107941 | Hypothetical protein |
| SaGU1_163 | CDS | + | 107941 | 108210 | Hypothetical protein |
| SaGU1_164 | CDS | + | 108295 | 108474 | Hypothetical protein |
| SaGU1_165 | CDS | - | 109074 | 108784 | Hypothetical protein |
| SaGU1_166 | CDS | - | 109412 | 109164 | Hypothetical protein |
| SaGU1_167 | CDS | - | 109671 | 109426 | Hypothetical protein |
| SaGU1_168 | CDS | - | 110094 | 109684 | Hypothetical protein |
| SaGU1_169 | CDS | - | 110648 | 110094 | Hypothetical protein |
| SaGU1_170 | CDS | - | 110895 | 110641 | Hypothetical protein |
| SaGU1_171 | CDS | - | 111086 | 110895 | Hypothetical protein |
| SaGU1_172 | CDS | - | 111568 | 111083 | Hypothetical protein |
| SaGU1_173 | CDS | - | 111815 | 111561 | Hypothetical protein |
| SaGU1_174 | CDS | - | 112273 | 111815 | Hypothetical protein |
| SaGU1_175 | CDS | - | 112710 | 112279 | Hypothetical protein |
| SaGU1_176 | CDS | - | 113220 | 112726 | Hypothetical protein |
| SaGU1_177 | CDS | - | 113637 | 113233 | Hypothetical protein |
| SaGU1_178 | CDS | - | 114341 | 113640 | Serine/threonine protein phosphatase |
| trna1 | tRNA | - | 114850 | 114779 | tRNA-Met-CAT |
| SaGU1_179 | CDS | - | 115678 | 115130 | Hypothetical protein |
| SaGU1_180 | CDS | - | 115900 | 115682 | Hypothetical protein |
| SaGU1_181 | CDS | - | 116095 | 115901 | Hypothetical protein |
| SaGU1_182 | CDS | - | 116822 | 116085 | Hypothetical protein |
| SaGU1_183 | CDS | - | 116989 | 116885 | Hypothetical protein |
| SaGU1_184 | CDS | - | 117240 | 117001 | Hypothetical protein |
| SaGU1_185 | CDS | - | 117628 | 117242 | Hypothetical protein |
| SaGU1_186 | CDS | - | 117898 | 117725 | Hypothetical protein |

| | | | | | |
|---|---|---|---|---|---|
| SaGU1_187 | CDS | - | 118421 | 117939 | Hypothetical protein |
| SaGU1_188 | CDS | - | 119013 | 118471 | Hypothetical protein |
| SaGU1_189 | CDS | - | 119543 | 119013 | Hypothetical protein |
| SaGU1_190 | CDS | - | 119710 | 119546 | Putative membrane protein |
| SaGU1_191 | CDS | - | 119991 | 119713 | Putative membrane protein |
| SaGU1_192 | CDS | - | 120836 | 119991 | Hypothetical protein |
| SaGU1_193 | CDS | - | 121967 | 120849 | AAA family ATPase |
| SaGU1_194 | CDS | - | 122446 | 122120 | Hypothetical protein |
| SaGU1_195 | CDS | - | 122855 | 122439 | Hypothetical protein |
| SaGU1_196 | CDS | - | 123290 | 122988 | Phage DNA-binding protein |
| SaGU1_197 | CDS | - | 123478 | 123290 | Hypothetical protein |
| SaGU1_198 | CDS | - | 123683 | 123522 | Hypothetical protein |
| SaGU1_199 | CDS | - | 125735 | 123684 | Hypothetical protein |
| SaGU1_200 | CDS | - | 126078 | 125815 | Hypothetical protein |
| SaGU1_201 | CDS | - | 126268 | 126095 | Hypothetical protein |
| SaGU1_202 | CDS | - | 126853 | 126275 | Hypothetical protein |
| SaGU1_203 | CDS | - | 127472 | 126846 | Nucleoside 2-deoxyribosyltransferase |
| SaGU1_204 | CDS | - | 128358 | 127465 | Putative DNA ligase |
| SaGU1_205 | CDS | - | 128586 | 128362 | Hypothetical protein |
| SaGU1_206 | CDS | - | 129394 | 128654 | Phosphate starvation-inducible protein PhoH, predicted ATPase |
| SaGU1_207 | CDS | - | 130060 | 129446 | Hypothetical protein |
| SaGU1_208 | CDS | - | 130501 | 130076 | Phage ribonuclease H |
| SaGU1_209 | CDS | - | 130682 | 130491 | Hypothetical protein |
| SaGU1_210 | CDS | - | 131346 | 130705 | Hypothetical protein |
| SaGU1_211 | CDS | - | 131566 | 131336 | Hypothetical protein |
| SaGU1_212 | CDS | - | 131796 | 131569 | Hypothetical protein |
| SaGU1_213 | CDS | - | 132597 | 131905 | Immunodominant staphylococcal antigen A precursor |
| SaGU1_214 | CDS | - | 133588 | 132794 | Putative membrane protein |
| SaGU1_215 | CDS | - | 133896 | 133588 | Hypothetical protein |
| SaGU1_216 | CDS | - | 135498 | 134011 | Putative lysin |
| SaGU1_217 | CDS | - | 136001 | 135498 | Putative holin |
| SaGU1_218 | CDS | - | 136271 | 136086 | Hypothetical protein |
| trna2 | tRNA | - | 136502 | 136431 | tRNA-pseudo-CCA |
| trna3 | tRNA | - | 136581 | 136509 | tRNA-Phe-GAA |
| trna4 | tRNA | - | 136661 | 136588 | tRNA-Asp-GTC |
| SaGU1_219 | CDS | - | 138032 | 137814 | Hypothetical protein |
| SaGU1_220 | CDS | - | 138722 | 138513 | Hypothetical protein |
| SaGU1_221 | CDS | - | 139067 | 138735 | Hypothetical protein |
| SaGU1_222 | CDS | - | 139406 | 139080 | Hypothetical protein |
| SaGU1_223 | CDS | + | 139846 | 140232 | Putative membrane protein |
| SaGU1_224 | CDS | + | 140210 | 140488 | Hypothetical protein |
| SaGU1_225 | CDS | + | 140485 | 140895 | Hypothetical protein |
